# SCENIC: Single-cell regulatory network inference and clustering

**DOI:** 10.1101/144501

**Authors:** Sara Aibar, Carmen Bravo González-Blas, Thomas Moerman, Jasper Wouters, Vân Anh Huynh-Thu, Hana Imrichova, Zeynep Kalender Atak, Gert Hulselmans, Michael Dewaele, Florian Rambow, Pierre Geurts, Jan Aerts, Jean-Christophe Marine, Joost van den Oord, Stein Aerts

## Abstract

Single-cell RNA-seq allows building cell atlases of any given tissue and infer the dynamics of cellular state transitions during developmental or disease trajectories. Both the maintenance and transitions of cell states are encoded by regulatory programs in the genome sequence. However, this regulatory code has not yet been exploited to guide the identification of cellular states from single-cell RNA-seq data. Here we describe a computational resource, called SCENIC (Single Cell rEgulatory Network Inference and Clustering), for the simultaneous reconstruction of gene regulatory networks (GRNs) and the identification of stable cell states, using single-cell RNA-seq data. SCENIC outperforms existing approaches at the level of cell clustering and transcription factor identification. Importantly, we show that cell state identification based on GRNs is robust towards batch-effects and technical-biases. We applied SCENIC to a compendium of single-cell data from the mouse and human brain and demonstrate that the proper combinations of transcription factors, target genes, enhancers, and cell types can be identified. Moreover, we used SCENIC to map the cell state landscape in melanoma and identified a gene regulatory network underlying a proliferative melanoma state driven by MITF and STAT and a contrasting network controlling an invasive state governed by NFATC2 and NFIB. We further validated these predictions by showing that two transcription factors are predominantly expressed in early metastatic sentinel lymph nodes. In summary, SCENIC is the first method to analyze scRNA-seq data using a network-centric, rather than cell-centric approach. SCENIC is generic, easy to use, and flexible, and allows for the simultaneous tracing of genomic regulatory programs and the mapping of cellular identities emerging from these programs. Availability: SCENIC is available as an R workflow based on three new R/Bioconductor packages: *GENIE3, RcisTarget* and *AUCell.* As scalable alternative to GENIE3, we also provide *GRNboost,* paving the way towards the network analysis across millions of single cells.

## Introduction

The transcriptional state of a cell emerges from an underlying gene regulatory network in which a limited number of transcription factors and co-factors regulate each other and their downstream target genes ^1^. Recent advances in single-cell transcriptome profiling have provided exciting opportunities for a high-resolution identification and clustering of transcriptional states, and to identify trajectories of transitions between states, for example during differentiation ^2,3^. Statistical techniques and bioinformatics methods have been optimized for single-cell RNA-seq ^4^, including methods for expression normalization ^5,6^, differential expression analysis ^7–12^, clustering ^13–16^, dimensionality reduction ^17,18^, trajectory inference ^19^, and rare cell type identification^17,20^. Although these methods have led to significant new biological insights, it is still unclear whether specific and robust GRNs underlying thus predicted cell states can be established. This may indeed be challenging given that at the single cell level, gene expression may, at least in part, be disconnected from the dynamics of transcription factor inputs due to stochastic variation of gene expression consecutive to, for example, transcriptional bursting ^21–23^. In addition, single-cell approaches usually yield low detection coverage of expressed genes and high numbers of dropouts ^3,4^. Linking the genomic regulatory code to single-cell gene expression variation, however, may allow exploiting the regulatory genome to optimize the analysis of single-cell RNA-seq data, to overcome drop-outs and technical variation, and to guide the discovery and characterization of cellular states. Consistently, methods that exploit co-expression or networks for the analysis of single-cell RNA-seq data such as “network synthesis toolkit” ^24^, Pina’s approach ^25^, PAGODA ^13^, and SINCERA ^26^ have tentatively been developed. However, these do not make use of regulatory sequence analysis to predict interactions between transcription factors and target genes.

Here we developed a new method, called SCENIC (Single-Cell rEgulatory Network Inference and Clustering), to characterize gene regulatory networks using single-cell RNA-seq data, and simultaneously optimize cellular state identification using the inferred networks. We benchmarked each of the features of SCENIC against alternative approaches, and applied SCENIC on a variety of recently published single-cell RNA-seq data sets, covering different species (human, mouse), tissues (brain, retina, tumors), and biological (differentiation) or pathological (intratumor heterogeneity) processes. Our results show that GRNs constitute robust guides to identify high-resolution cellular states, and that single-cell RNA-seq data are well-suited to trace gene regulatory programs in which specific combinations of transcription factors drive cell type-specific transcriptomes.

## Results

### Simultaneous discovery of gene regulatory networks and cellular states in the mouse brain

We developed **SCENIC** (Single-Cell rEgulatory Network Inference and Clustering) to map gene regulatory networks from single-cell RNA-seq data, using a combination of co-expression network inference, transcription factor motif analysis, and network-based prediction of cellular subpopulations (Figure 1). All three steps are based on new R/Bioconductor packages (see Methods). To test SCENIC performances we applied it to a scRNA-seq data set with well-known cell types from the adult mouse brain previously described in Zeisel et al. ^16^. This data set has been used extensively for benchmarking purposes ^13,14,20,27–31^ and contains the main cell types in hippocampus and somatosensory cortex, namely neurons (pyramidal excitatory neurons, and interneurons), glia (astrocytes, oligodendrocytes, microglia), and endothelial cells. In the first step (Figure 1a), we inferred co-expression modules using an improved implementation of GENIE3 ^32^, the top-performing method for network inference in the DREAM challenge ^33^. GENIE3 identified coexpression modules associated to 1046 TFs, ranging in size from 52 to 12082 target genes (we use various thresholds for N, see Methods). Since GENIE3 uses Random Forest regression, it has the added value of allowing complex (e.g., non-linear) co-expression relationships between a TF and its candidate targets.

**Figure 1.**
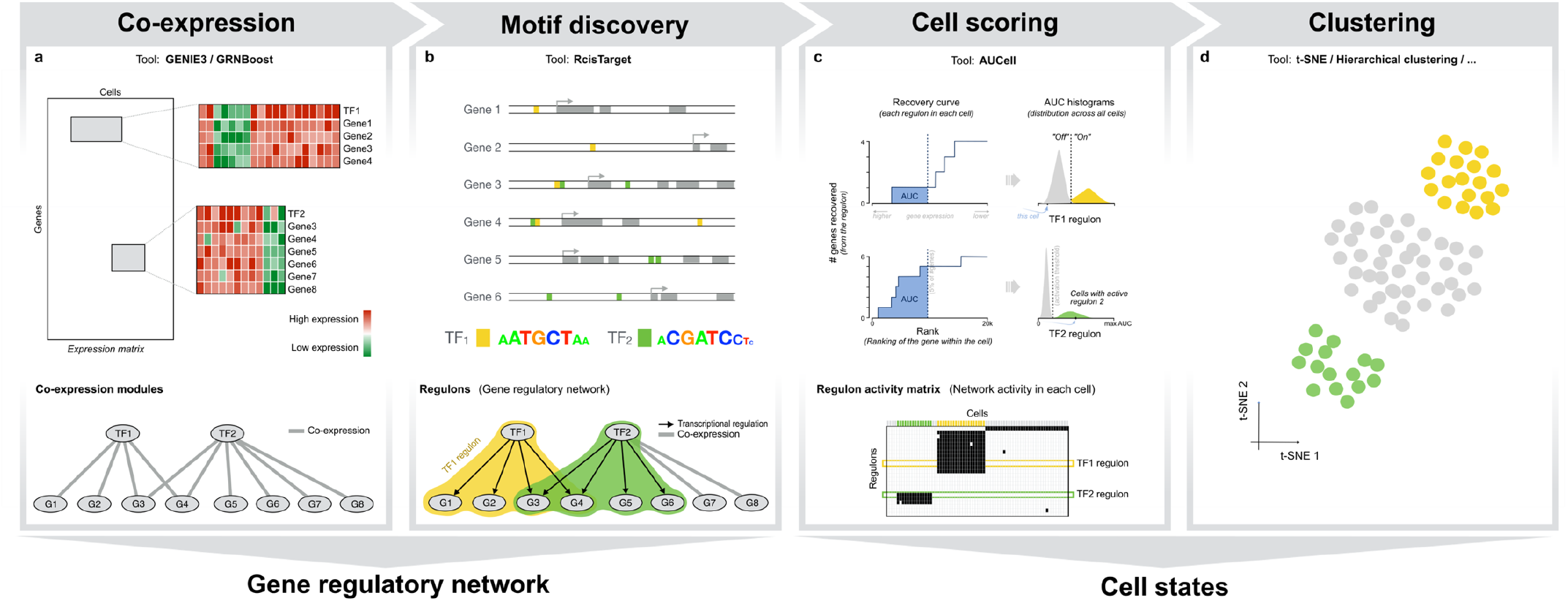
The SCENIC workflow. (**a**) In the first step, co-expression modules between transcription factors and candidate target genes are inferred with GENIE3 or GRNboost. Each module consists of a transcription factor together with its predicted targets, purely based on coexpression. (**b**) In the second step, each co-expressed module is analyzed with RcisTarget to identify enriched motifs; only modules and targets for which the motif of the TF is enriched are retained. Each TF together with its potential direct targets is a *regulon.* (**c**) In the third step, the activity of each regulon in each cell is evaluated using AUCell, which calculates the Area Under the recovery Curve. The AUCell scores are used to generate the Regulon Activity Matrix. This matrix can be binarized by setting an AUC threshold for each regulon, which will determine in which cells the regulon is “active”. (**d**) The Regulon Activity Matrix can be used to cluster the cells (e.g. t-SNE) and, thereby, identify cell types and states based on the shared activity of a regulatory subnetwork.

These co-expression modules contained many false positive predictions, and a mixture of direct and indirect interactions ^33^. Therefore, the second step of SCENIC analyzes all the co-expression modules using *cis*-regulatory motif analyses (Figure 1b). We reasoned that, if the DNA recognition motif of the regulator is significantly enriched in the upstream and intronic sequences of the coexpressed gene set, then this module is of higher confidence. Furthermore, for the selected modules we determined the optimal subset of direct targets, thereby pruning the initial module to remove indirect targets. For this motif enrichment step we used a new R implementation of i-cisTarget ^34^, called RcisTarget. Briefly, RcisTarget tests for enrichment of more than 18 thousand position weight matrices using a cross-species ranking approach (see Methods). On the mouse brain, RcisTarget identified 151 of the initial 1046 GENIE3 modules as significantly enriched for the motif of the co-expressed transcription factor (7% of the initial TFs). Next, RcisTarget determines for each enriched motif the optimal “leading edge” of direct targets. This results in a subset of pruned modules, with only putative direct targets, which we call *regulons.* The other modules, without support for direct regulation, are removed.

In the third step, SCENIC scores all cells for the activity of each regulon. This scoring is done with a new metric, called *AUCell* (Area Under a Cell), which is inspired by order statistics and recovery analyses (see Figure 1c and Methods). *AUCell* calculates the enrichment of the regulon as an area under the recovery curve (AUC) across the ranking of all genes in a particular cell, whereby genes are ranked by their expression value. This method is therefore independent of the gene expression units and the normalization procedure. Using the distribution of *AUCell* scores across all cells, an optimal subpopulation is selected (Figure 2a, S1). We validated AUCell using previously published neuronal and glial gene signatures, and found it more robust than using the mean of the normalized expression values across the gene signature (Figure S2).

**Figure 2.**
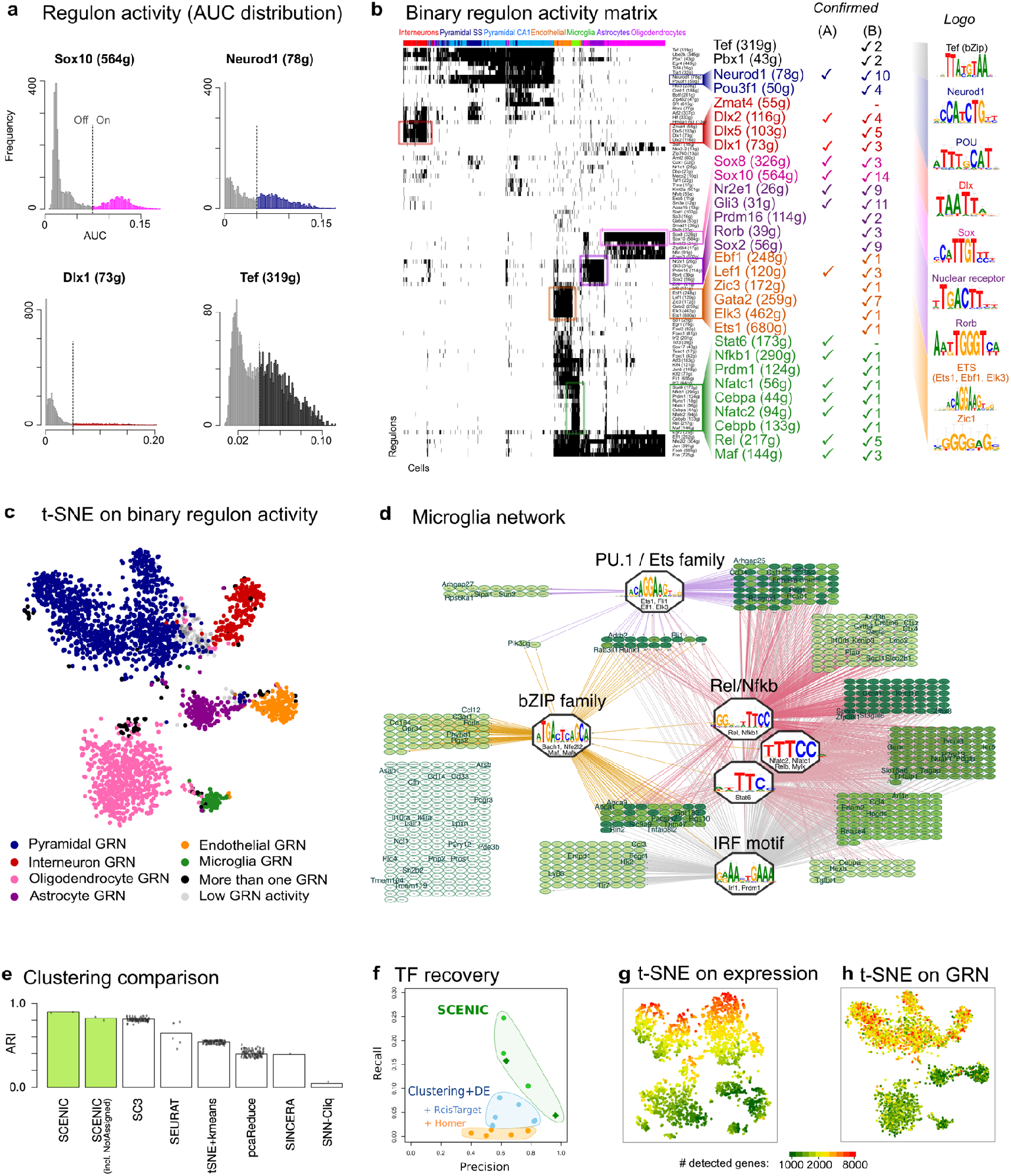
SCENIC analysis of the mouse brain. (**a**) AUC histograms for a few key regulons. The AUC allows to split the populations of cells with high versus low activity of a regulon. (**b**) The clustered regulon activity matrix reveals that the known cell types have distinct regulatory networks. The color bar above the heatmap indicates the cell type assigned by the authors of the dataset. Key regulons (rows) are magnified and colored according to the cell type in which they are active. The two annotation columns on the right indicate whether the TF is known to be relevant (A: for the cell type, manually curated from literature; B: brain-related TFs annotated by MGI), and the main DNA motifs. (**c**) t-SNE on the binary regulon activity matrix. Each cell is assigned the color of the most active regulons. (**d**) Microglia gene regulatory network. The regulons associated to microglia can be summarized based on the binding motif of the associated TF (network built in iRegulon). The genes that are included in a previously published microglia signature (Lavin et al. ^105^) are indicated by a larger font size; the color of the node indicates the number of regulators (lighter: fewer, darker: more). (**e**) Performance of different clustering methods on this dataset (ARI: Adjusted Rand Index, calculated taking as reference the cell-type assigned by the authors). (**f**) Comparison of the precision and recall of TF prediction by SCENIC versus other methods. (**g**) t-SNE on the expression matrix (same input as to SCENIC: UMI counts with no further normalization) and (**h**) t-SNE on the binary regulon activity matrix. Both t-SNEs are PCA-based and colored according to the number of genes detected (expression over 0) in each cell.

When applied to the 151 regulons inferred on the mouse brain data, AUCell leads to a binary regulon activity matrix (Figure 1c, Figure 2a-b). Hierarchical clustering of this matrix uncovered the expected cell types –which are in full agreement with the cell types annotated by the authors of this study–, thus demonstrating that the activity of gene regulatory subnetworks can accurately predict cell types (Figure 2b-c). This is not the case when only gene expression of the predicted transcription factors is used for clustering, indicating that the target genes provide robustness (Figure S3). Moreover, the regulons and the top transcription factor predictions, were consistent with their previously established roles in the respective cell types: Mef2c, Neurod, Creb, and Egr4 in neurons; Sox10 in oligodendrocytes; Dlx1/2 in interneurons; and Lef1 in endothelial cells. Particularly, the predicted network of microglia contains many well-known regulators of microglial fate and/or microglial activation, including PU.1, Nfkb, Irf, and AP-1/Maf (Figure 2b, 2d). When we compared the predicted microglial network to previously published gene signatures of microglial “activation” in a mouse CK-p25 Alzheimer’s disease (AD) model ^35^, we found the microglia network to be strongly activated and the neuronal network to be down-regulated during AD progression (Figure S4), indicating that the microglia network captures a relevant regulatory program.

SCENIC therefore successfully (1) identifies TF-driven regulatory modules based on co-expression; (2) trims these modules towards direct TF-target relationships by motif discovery; and (3) maps the activity of the array of regulons onto single cells, thereby generating a regulon activity matrix, which allows accurate identification of cellular states. The network-driven single-cell clustering provides robust cell clustering, alongside meaningful biological insights into master regulators and gene regulatory networks underlying specific cell types.

### Comparison with other methods

To further evaluate the performance of SCENIC, we performed four benchmark comparisons, each assessing a different aspect of SCENIC: (1) cell clustering accuracy; (2) TF identification; (3) automatic normalization; and (4) automatic batch effect removal. In this section, we present the first three comparisons using the mouse brain data, while the detection of batch effects will be discussed in sections below using a cross-species analysis, and using two related cancer data sets.

To compare the accuracy of cell clustering based on SCENIC GRNs with standard clustering methods, we used the recent benchmark assay from the SC3 publication ^14^, based on the adjusted Rand index (ARI) ^36^. The ARI determines whether pairs of cell are correctly assigned to the same cluster, using as ground truth the clustering and annotation provided by the authors of the original study. SCENIC clusters were highly reliable, with an overall sensitivity of 0.88 and specificity of 0.99, and a clustering performance at least as good as the best performing method thus far (ARI > 0.80), namely SC3, while outcompeting all other methods (Figure 2e). To further assess the robustness of cell clustering by SCENIC, we re-analyzed the mouse brain data, now either using only 100 randomly selected cells (to simulate small data sets), or using 1/3 of the sequencing reads (to simulate low-coverage data sets). Interestingly, SCENIC identified cell types that are represented by only few cells (e.g. 2-6 cells from microglia, astrocytes or interneurons) and is relatively robust to drop-outs (Figure S5).

In the second comparison, we tested the performance of SCENIC in identifying correct transcription factors. To do so, we first created a set of 571 “true positive” brain-related TFs based on annotation from the Mouse Genome Informatics database (MGI) and GO ^38,39^ (see Methods), which we used to calculate the recall and precision of TF identification. We compared SCENIC with a “standard” analysis pipeline, consisting of clustering, differential expression (DE) analysis, and motif discovery. Applying various clustering and DE methods and parameters (see Methods) led to multiple gene signatures per cell type, which were analyzed for enriched motifs using Homer (we chose Homer because it was the second-best performer in the iRegulon benchmark ^40^). This analysis showed that SCENIC significantly outperforms standard analyses approaches (Figure 2f). For example, at >0.95 precision, SCENIC finds 25 relevant TFs, while the next performing method detected only 10 relevant TFs at even lower precision (~0.85). Overall, SCENIC could potentially identify up to 212 out of 310 TFs detected at protein level in the mouse brain by Zhou et al. (65 TFs would be missed because they are not detected in the scRNA-seq dataset, and only 38 because they have no motifs annotated in RcisTarget databases, Figure S6b).

In the third comparison, we investigated how gene expression normalization impacts gene regulatory network inference. Applying SCENIC to raw data versus normalized data gave similar clustering accuracies, differing only 3% in ARI score (0.87-0.90), and concordant TF recoveries, (26/30 TFs are common). Related to normalization, we found an interesting difference between the t-SNE plots based on the raw expression matrix versus the regulon activity matrix. While the clustering applied directly on the expression matrix is strongly biased towards the number of genes expressed in a cell, the clustering based on SCENIC effectively corrects for the intra-cluster bias, while the true biological difference between neurons (more genes expressed) and glia (less genes expressed) is unaffected ^41,42^ (Figure 2g-h). Thus, SCENIC removes the intra-cluster bias without having to decide which normalization method to use.

In conclusion, SCENIC competes with the best clustering methods to discovering cell types and correctly assigning cells to each cell type; but SCENIC goes beyond existing methods by reducing data dimensionality using TF regulons rather than principal components, thereby accounting for noise and removing technical biases, and uncovering master regulators and gene regulatory networks for each cell type.

### Inferring cross-species cell states and networks

Our analysis of the mouse brain revealed a single cluster of interneurons, with an underlying network driven by the highly significant factors Dlx1 and Dlx2 (Figure 2b, 2c). To validate this Dlx1/2 network, we analyzed a recently published single-nuclei RNA-seq data set of the human brain by Lake et al ^43^. On the human data, SCENIC identifies a cluster of interneurons strongly driven by DLX1/2 (see Figure S7 for the entire regulon activity matrix), having the same recognition motif as in mouse. Comparing the predicted target genes of mouse Dlx1/2 and human DLX1/2 revealed a conserved set of targets including DLX1 itself (auto-regulatory), NR2E1, SP8, SOX1, NXPH1, and IGF1, which are all closely associated to DLX1/2 in terms of protein-protein interactions or literature evidence (Figure 3a-b). This cross-species analysis cross-validated the predicted DLX1/2 targets in interneurons, and yielded a core set of conserved targets, alongside a subset of species-specific target genes. In addition, the regulon-based clustering of the Lake et al. human nuclei featured three subtypes of interneurons, corresponding to the known interneuron subtypes (PVALB, VIP, SST, see Figure S8). This contrasts with the results for the mouse dataset, where the interneurons make up one homogeneous cluster of cells without clear subtypes, due to the experimental setup of Zeisel et al. dataset, where only one type of interneurons (Htr3a positive cells) were purified.

**Figure 3.**
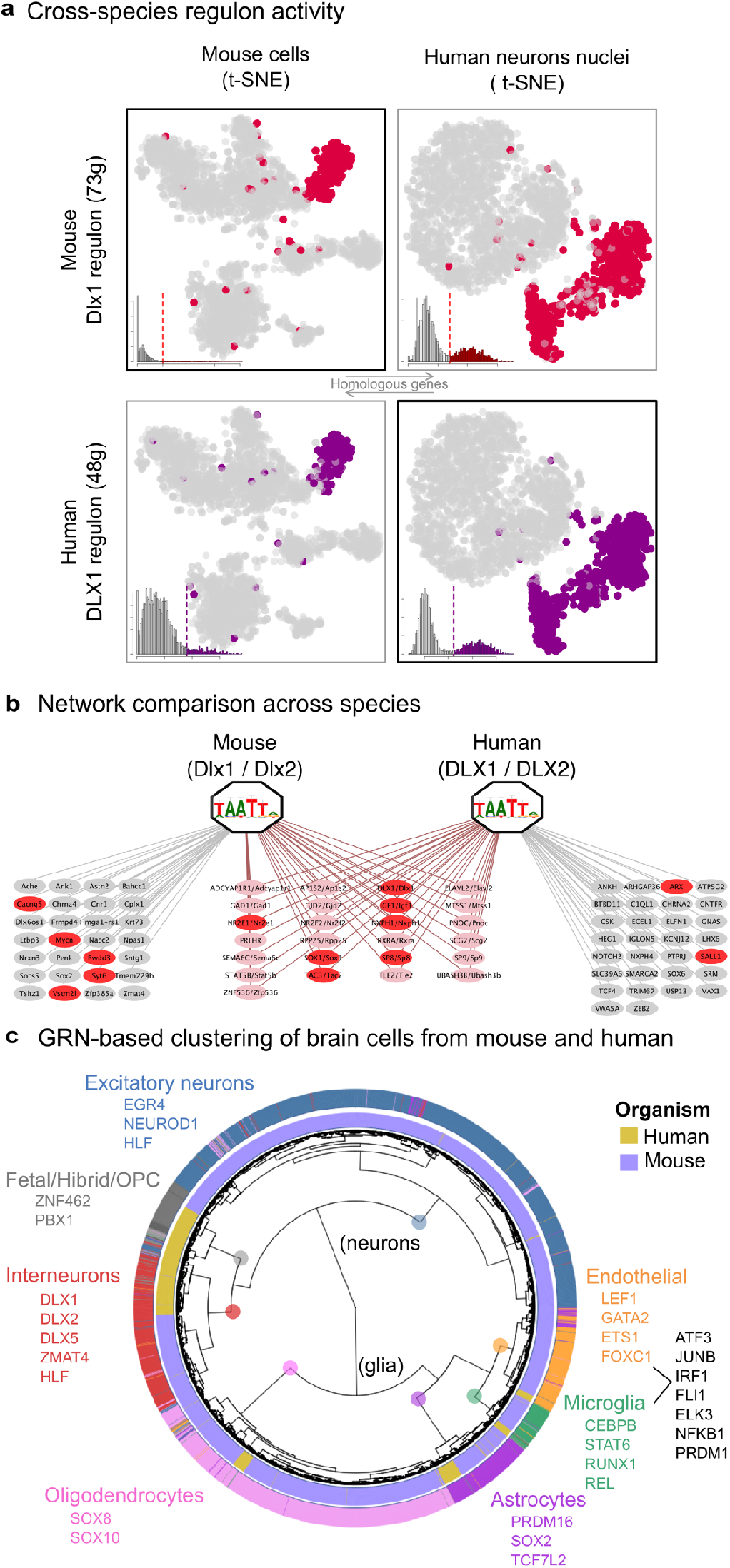
Cross-species comparison of networks and cell types. (**a**) Reciprocal activity of human and mouse Dlx1/2 regulons on mouse and human single-cell data. In both cases, the Dlx network is mainly active in interneurons, indicating strong agreement between the species. (**b**) Shared targets between DLX1/2 regulons inferred from mouse and human. The genes highlighted in red also have associations with Dlx1/2 in GeneMANIA ^106^ (protein-protein interactions, genetic interactions, co-expression, or literature co-mentioning). (**c**) Clustering based on GRN activity allows combining cells from human and mouse: The clustering is driven by the cell type, rather than by the organism (unlike the clustering resulting from the normalized expression matrix, Figure S9). Colored TF names show conserved regulons between human and mouse.

Encouraged by this finding, we asked whether cross-species network activity could be exploited more generally to analyze cross-species scRNA-seq, possibly overcoming “batch effects” that are notorious in cross-species analysis ^44^. Analogous to the cross-species DLX validation, we used SCENIC to “train” regulons from one species, and then score cells of another species with the orthologous genes using AUCell, and vice versa. Note that since AUCell is independent of the units of the dataset (as it is a rank-based approach), it was possible to apply it to simultaneously study mouse and human brain^41^ data. In contrast to standard clustering based on normalized expression, which yields a strong species-driven clustering (Figure S9), the SCENIC analysis effectively grouped cells by cell type first, and then by species, as expected (i.e., a human and mouse interneuron are more similar to each other than a human interneuron and a human oligodendrocyte) (Figure 3c). Interestingly, SCENIC identified an “interneuron-like” and a “excitatory neuron-like” subpopulation within the fetal quiescent cells in the human data set, expressing DLX1,2,5 and MAF, and NEUROD1.

Together, the analysis of Lake et al. dataset confirmed that *single-nuclei* RNA-seq are also amenable to gene regulatory network analysis, and provided putative regulators for interneuron subtypes. By comparing mouse and human single-cell data sets, we confirmed that some gene regulatory networks are conserved across species, and validated a predicted DLX network across interneurons. Finally, we provide an example of how regulatory network analysis can overcome species-specific biases.

### Upscaling to big data

With new developments in single-cell RNA-seq technologies, including the use of droplet microfluidics ^45,46^ and the release of commercial platforms such as 10X Genomics, datasets with increasingly large numbers of single cells are becoming available. A recent demonstration from 10X Genomics released a dataset containing 1.3 million cells from the embryonic mouse brain ^47^ (E18.5 hippocampus, cortex, and ventricular zone). To apply network inference on such large datasets we suggest two complementary approaches. In the first approach, we infer the GRN from a sub-sampled data set, and re-include all cells in the third step, for the AUCell scoring. We tested this approach using GENIE3 on a large Drop-seq dataset containing 44808 single cells from the mouse retina from Macosko et al. ^45^. For GENIE3 we selected the same 11020 cells as the authors used (the authors made that selection based on the number of genes expressed, being at least 900 genes ^45^, see Methods) and derived 123 regulons. The identified master regulators, such as Sox8/9, Hes1, Rax, Nr1h4, Srf, and Nr2e1 for the Muller glia ^48–51^, illustrated that correct networks can be inferred even on sparse data. All 44K cells were then scored with these regulons to build the full t-SNE plot (Figure S10). In the second approach, we solved the computational efficiency problem by using more efficient machine learning and big data handling solutions. Particularly, we implemented a new variant of GENIE3 in Scala that replaces the Random Forest regression with gradient boosting. This new implementation drastically reduces the time needed to infer a GRN. For example, network inference on the mouse brain data (3K cells) is at least 30 times faster than GENIE3 on a single node, while retaining the accuracy of TF and target prediction (Figure S11a). Additionally, we exploited the “embarrassingly parallel” nature of this inference method (the regulators are predicted individually for each target), designing GRNboost as a distributed algorithm, and using Apache Spark ^52^ to coordinate the computations across a compute cluster. We applied GRNboost on 3K, 10K, and 100K cells from the embryonic mouse brain, and were able to infer gene regulatory networks within 18 and 37 minutes, and 8 hours respectively (Figure S11b). Using two technical replicates of 100K cells, we found that target gene predictions by GRNBoost were robust (Figure S12). This method will pave the way to network inference on very large data sets, such as the soon available Human Cell Atlas ^53^.

### SCENIC unravels the heterogeneity of regulatory states in tumours

Understanding the mechanisms driving intratumor heterogeneity has become a priority in cancer biology as, among others, it represents an essential step towards the development of improved rational combination treatments ^54^. Single-cell RNA-seq provides unprecedented opportunities to gain insights into the molecular determinants of cancer cell phenotypic diversity, and mapping the gene regulatory networks may significantly contribute to this. However due to tumor-specific mutations and complex genomic aberrations, the identification of cancer cell states is more challenging than characterizing normal cell types ^55^. To further test SCENIC utility for the analysis of such complex datasets, we run it on two recently published scRNA-seq data sets, the first obtained from six oligodendroglioma tumours ^56^ (4043 cancer cells), and the second from fourteen melanoma lesions ^55^ (1252 cancer cells). Standard clustering of the oligodendroglioma expression matrix resulted in clusters of cancer cells by the tumour of origin, similar to previous observations ^55,57^ (Figure 4a), due to strong inter-tumor heterogeneity ^54^. Critically, SCENIC-based cell clustering revealed three cancer cell states shared across tumours namely “astrocyte-like”, “oligodendrocyte-like”, and cancer stem cells (including cycling cells). Each state is driven by expected/relevant TFs, including SOX10/4/8, OLIG1/2, and ASCL1 for the oligodendrocyte-like state, and SOX9, NFIB, AP-1 for the astrocyte-like state, and E2F, FOXM1 for the cycling cells. We further validated the cluster of cycling cells by comparing to the author’s labels and by examining gene signatures related to the cell cycle using GO-terms (Figure S13). Interestingly, when we used diffusion plots with the binary matrix (Figure 4b) rather than t-SNE, SCENIC was able to reconstruct the differentiation trajectory (from stem-like, undifferentiated, to the oligodendrocyte-like branch and astrocyte-like branch). Note that this cancer trajectory represents a different “trajectory” compared to normal oligodendrocyte differentiation (Figure S14 for the SCENIC analysis of 5069 oligodendrocytes ^58^). We then asked to what extent these three clusters can be identified by other methods (Figure 4a). This revealed that SCENIC can compete with dedicated batch-effect removal methods, such as Combat ^59,60^ and Limma ^61,62^, when used to explicitly remove the tumor-batch effect. With SCENIC, this batch-effect is removed automatically, without requiring prior information about the source of the batch effect.

**Figure 4.**
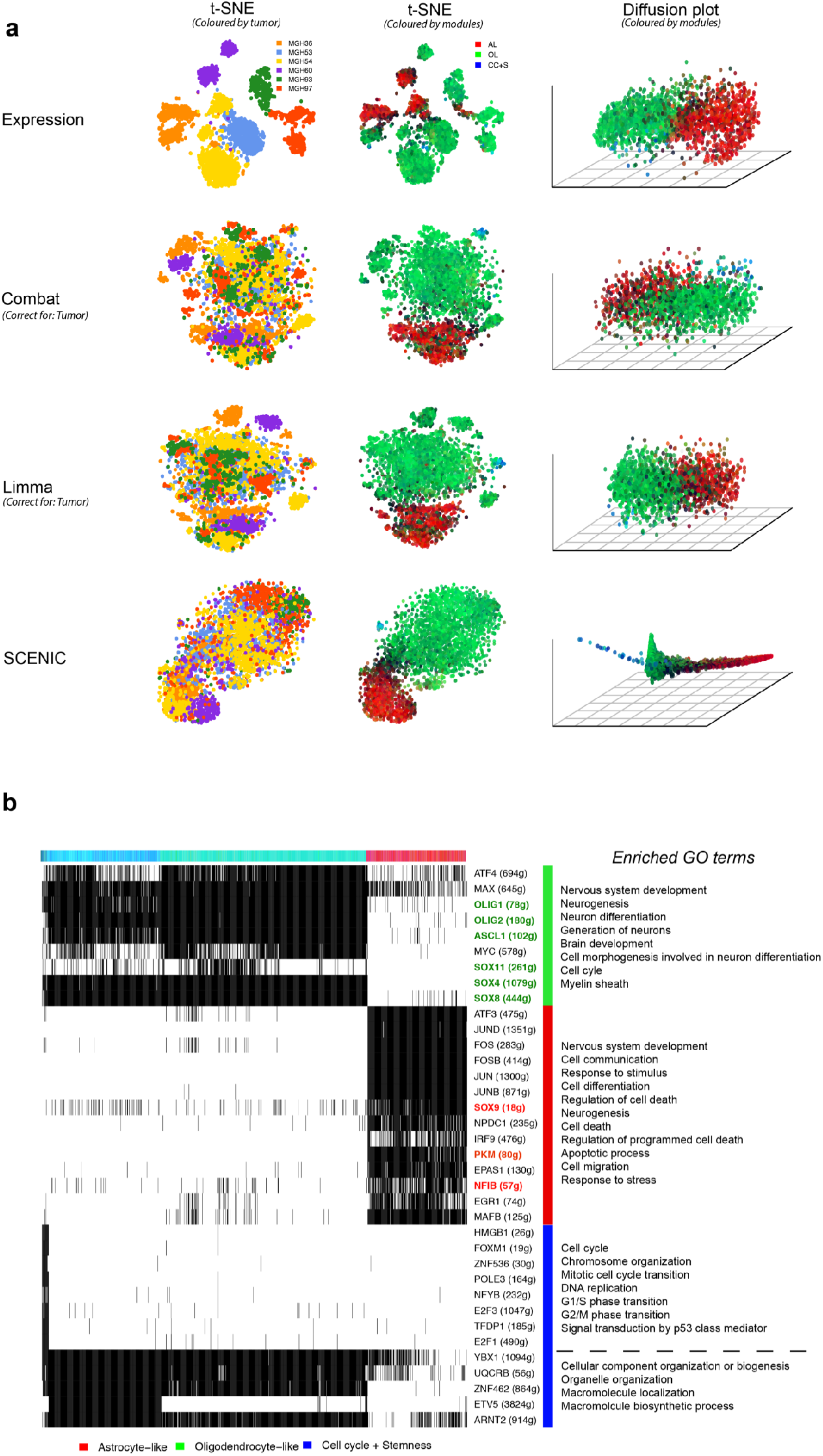
SCENIC overcomes tumor batch effect and recovers relevant cell types and GRNs in oligodendroglioma. (**a**) Comparison of batch-effect removal methods on an oligodendroglioma dataset. t-SNEs and diffusion plots on the raw expression matrix (first row), after correcting by tumor of origin with Combat or Limma (rows 2-3), or on the binary activity matrix from SCENIC (row 4). The cells are coloured based on the tumor of origin or GRN activity (red: astrocyte-like regulons, green: oligodendrocyte-like regulons, blue: regulons related to cell cycle or stemness). (**b**) Simplified binary regulon activity matrix (output of SCENIC) for the oligodendroglioma dataset. Highlighted regulons (colored TF names) are known to be characteristic in oligodendrocytes or astrocytes, respectively.

We observed a similar batch effect correction on the melanoma data, where SCENIC clusters cancer cells from different tumors together, in contrast to the standard clustering (Figure 5a-c). Similar to oligodendroglioma, we identified a cluster of cycling cells, driven by similar transcription factors (e.g., E2F1/2/8 and MYBL2, Figure 5f-g). The targets of these factors are strongly enriched for G1/S phase transition according to pathway enrichment analysis (Figure 5h). We validated this state by comparing gene expression across all cells using the gene signatures derived from cell cycle related GO terms (Figure 5n-o).

**Figure 5.**
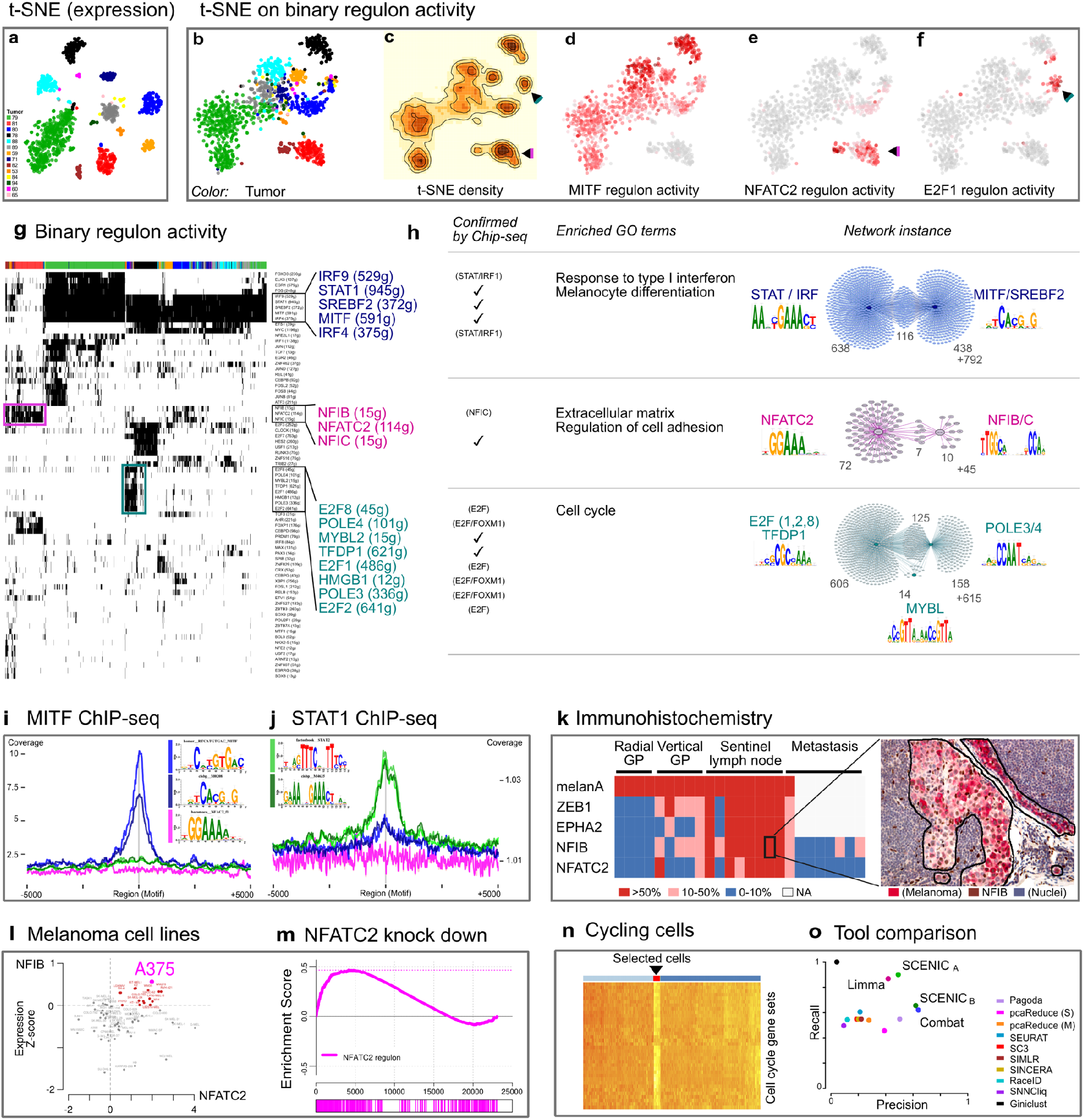
SCENIC reveals melanoma heterogeneity. (**a**) t-SNE on the raw melanoma expression matrix colored by tumor of origin. (**b-f**) t-SNE on the binary activity matrix after applying SCENIC. (b: cells colored by tumor of origin, c: density plot, d: cells colored by the AUC of the MITF regulon, **e**,f: same for NFATC2 and E2F1). (**g**) Binary regulon activity matrix for this dataset. The color bar above the heatmap indicates the tumor of origin; regulons associated to the cell cycle (green), invasive (pink) and proliferative (blue) states are zoomed in. (**h**) Three most dominant networks: MITF^high^, usually known as proliferative, (MITF/STAT); MITF^low^, invasive state, (NFATC2/NFIB); and cell cycle (E2F). *Confirmed by ChIP-seq:* a tick indicates that the regulon presents enrichment of targets in a ChIP-seq dataset for the same transcription factor. (**i-j**) Aggregation plots for MITF and STAT1 ChIP-seq signal on the predicted target regions of MITF, STAT, and NFATC2, showing high specificity of the predictions. (**k**) Immunohistochemistry (IHC) on 25 human melanomas using NFATC2, NFIB, ZEB1, and EPHA2 antibodies. Biopsies are taken from primary lesions (9) in radial growth phase (RGP) or vertical growth phase (VGP), sentinel lymph nodes (8), and metastatic lesions (8). On the left, a heatmap summarizes the results of the IHC indicating the percentage of cells that are positive for each marker in the given sample. On the right, a representative example of IHC for NFIB on a sentinel lymph node (for additional images, see Figure S18; red, melanA, a marker for melanoma cells; brown, NFIB; blue, hematoxylin). (**l**) Z-score normalized expression of NFIB and NFATC2 across melanoma cell lines from COSMIC. (**m**) GSEA plot for genes differentially expressed after NFATC2 knock-down in the A375 cell line: the predicted NFATC2 targets are significantly up-regulated in the NFATC2 knock-down. (**n**) Cycling cells selected based on GO terms related to cell cycle. (**o**) Comparison of the capacity of different methods to identify the cycling cells (SCENIC_A_: High-confidence cells, SCENIC_B_: Lower confidence/regulon activity).

Interesting, the melanoma cells fall largely into two groups, one corresponding to a MITF^high^ state, showing also high *TYR* expression *(TYR* is a target gene of MITF [re]), and an MITF^low^ state (Figure 5d-e, Figure S15 and S16). For the MITF^high^ state, the two transcription factors with the most significant co-expression modules and the highest accompanying motif enrichment in these modules, are MITF and STAT. MITF is a known driver of (proliferative) melanoma ^63^, but its cooperativity with STAT in melanoma cells is a new prediction. To validate the predicted target genes for MITF and STAT1 we used previously published ChIP-seq data and found significant enrichment of ChIP-seq peaks at the predicted target regions (Figure 5h). Furthermore, these predictions show high specificity, since MITF ChIP-seq peaks are not enriched on predictions of other factors (Figure 5i-j).

The MITF^low^ cluster showed up-regulated WNT5A, LOXL2 and ZEB1 expression (both known markers of the invasive state ^64,65^), and correlates significantly with previously published invasive gene signatures (Figure S17). However, unlike the ‘classical’ invasive cell state, this MITF^low^ state retains SOX10 expression (Figure S16). The two top transcription factors in this state are NFATC2 (114 predicted target genes) and NFIB (15 predicted target genes). NFATC2 is involved in melanoma dedifferentiation to a stem cell fate and immune escape ^66^. NFIB on the other hand is linked to stem cell behavior of hair follicle and melanocyte stem cells ^67^ and plays an important role in metastatic progression of small cell lung cancer (SCLC) ^68^. To further explore the potential role of the new regulators NFATC2 and NFIB in the MITF^low^ melanoma state, we performed immunohistochemistry on 25 melanoma specimens with varying tumor progression, i.e., 9 primary melanomas (4 in radial growth phase and 5 in vertical growth phase), 8 melanoma-containing sentinel lymph nodes and 8 melanoma metastases. Interestingly, we found the highest NFIB and NFATC2 expression in the sentinel lymph nodes, co-localizing with ZEB1 expression, suggesting a relationship with the earliest metastatic events (Figure 5k and S16).

To further investigate the role of NFATC2 in this cell state we knocked-down NFATC2 using siRNA (see Methods) in a melanoma cell line that shows high NFATC2 and NFIB expression, namely A375 (Figure 5I), and performed RNA-seq to compare the transcriptome of A375 baseline with NFATC2 knock-down. The predicted NFATC2 target genes were significantly upregulated upon NFATC2 knockdown (Figure 5m). This is consistent with previously established role as a repressor ^69–71^. Interestingly, genes involved in regulation of cell adhesion and extracellular matrix and several previously-published gene signatures representing the melanoma invasive state are up-regulated (Table S1).

In conclusion, in addition to the well-described proliferative MITF^high^ melanoma state, SCENIC identified several new regulons for an in vivo MITF^low^ invasive melanoma state and novel regulators governing their respective transcriptional networks, which may play important roles in the progression of the disease.

## Discussion

We present a new computational approach that significantly improves the identification of cell types or cellular states present in a complex, heterogeneous biological sample and their respective master regulators and underlying GRNs. These are simultaneously determined using a combination of regulatory network inference, *cis*-regulatory motif enrichment, and identification of sub-populations based on rank statistics. For each of these three steps, we developed an R/Bioconductor package, accompanied by tutorials and integrative scripts that implement the SCENIC workflow.

### First network inference, then cell clustering

Standard analysis of single-cell RNA-seq data consists of data normalization, clustering cells into cell types, visualizing in t-SNE plots, and identification of gene signatures (marker genes) for each cell type ^4^. Here, we explored an alternative approach, to first analyze a single-cell RNA-seq expression matrix for gene co-expression relationships, yielding co-expressed gene modules. Compared to the multitude of methods that perform cell-centric analyses, only few perform gene-centric analyses, such as PAGODA ^13^. SCENIC goes beyond these methods by inferring *modules* of a transcription factor and its putative target genes. To achieve high-quality modules, we combined existing methods for TF-target inference that were originally developed for bulk RNA-seq or microarray data, namely GENIE3 ^32^ and gradient boosting ^72^, and found that they perform adequately on single-cell RNA-seq data. To overcome the excess of false-positive TF-target relations, we augmented these modules with cis-regulatory motif analysis, converting them into *regulons.* We found that such a regulon-centric approach provides several advantages. Firstly, it retains more transcription factors compared to a differential expression analysis. Secondly, the regulons are immune to drop-outs, because individual missing values are compensated by the network neighborhood, providing a more robust quantification of single-cell gene expression levels. By using the inferred regulons as features, we achieved a biological dimensionality reduction, yielding a regulon activity matrix, rather than a statistical transformation using PCA or similar approaches. Thirdly, clustering cells based on similar regulons is more accurate than most of the existing clustering algorithms, and is less sensitive to normalization. Clustering methods use a normalized gene expression matrix, which can be biased towards the number of genes detected in a cell, or the sample of origin. SCENIC effectively removes such technical biases by measuring the activity of subnetworks, rather than single genes.

There are still limitations to using transcription factor motifs to filter and prune co-expression modules, the most obvious being that not for all transcription factors motifs are available, that some factors have motifs with higher information content than others, and that not all transcription factors are co-expressed with their target genes.

### Finite network diversity in the brain

Applying SCENIC to the mouse and the human brain, we found that distinct cell states can be defined by stable, discrete configurations of a gene regulatory network in which specific combinations of “master” transcription factors regulate a critical subset of the genes expressed in that cell type. This finding reinforces the premise that a finite number of gene regulatory networks are encoded in the genome. The networks overall match almost perfectly with the known cell types in the brain, even up to the three known subpopulations (PV, SST, and VIP) of interneurons, which can be combined with other cellular states such as “stress/activation” or cell cycle. This is an interesting observation, because it suggests that the major cell types have in fact been discovered, that they can be confirmed by this unbiased approach of single-cell RNA-seq, and that they correspond to specific gene regulatory network configurations. On the other hand, this “unsophisticated view” on cell identity opens an intriguing question, namely how the identity of more specialized subpopulations of neurons and glia are controlled, since differences in gene regulatory networks are not obvious at the current resolution. A related question is whether transcriptional switches during cellular differentiation are controlled by different gene regulatory networks than the parental or child state. When considering the human oligodendroglioma and mouse oligodendrocyte data set, we found few dominant networks (e.g., stem cell, oligodendrocyte-like, and astrocyte-like). Transitions between these main states must occur rapidly, since only few cells were found in transition.

### Recurrent cancer gene networks compete with tumor-driven transcription

Single cells from cancer biopsies often cluster by their tumor/patient of origin (e.g. ^55–57,73^, Figure 3 and 6a). This contrasts with bulk tumor analyses, where different subtypes can be identified for most tumor types ^74^. The apparent differences between the tumors at single-cell level may be due to differences in copy number profiles, which are unique for each tumor and can have a strong impact on the gene expression profile ^55,57,75^. The use of gene regulatory networks effectively compensates for this effect, allowing the cells to be grouped by recurrent cancer cell states, thus more easily revealing the subtypes known from bulk analyses. The application of SCENIC to the oligodendroglioma and melanoma datasets clearly illustrates this possibility. Interestingly, our analysis revealed that most oligodendroglioma contain cells from both identified subtypes, namely oligodendrocyte-like and astrocyte-like, indicating that the tumor subtype identified through bulk analysis ^76^ is likely to reflect the proportions of cells adopting one of these cell states within a given tumor. In contrast, there was no obvious co-occurrence of the identified melanoma cell states, namely proliferative and invasive, within the same melanoma lesion. The only cell state shared between different tumours is defined as the cycling cells cluster, which strongly resembles the one identified in oligodendroglioma. Note that for oligodendroglioma, these cycling cells likely represent cancer stem cells, as validated by the authors ^75^. It is therefore tempting to speculate that this cycling subpopulation may also represent a melanoma stem cell population. However, further experiments would be needed to confirm this hypothesis.

SCENIC identified an “invasive” state largely shared by two of the 14 biopsies, both resected from auxiliary lymph nodes. This state, unlike the in vitro invasive state, which is driven by AP-1 and TEAD factors, this “in vivo” invasive state features distinct transcription factors, including NFATC2 and NFIB, which we confirmed to be expressed in early metastatic melanoma cells (i.e. in the initial, small tumors in the sentinel lymph node, by immunohistochemistry). Using gene expression analysis after NFATC2 knock down, we identified NFATC2 as a transcriptional repressor of the AP-1 target genes. Thus, these observations suggest that NFATC2 may act as a transcriptional break that cells need to overcome to switch to a full-blown invasive cell state. NFATC2 is itself a JUN target ^77^, and may constitute a negative feedback mechanism. A similar repressor function of NFATC2 has been previously observed in breast cancer ^78^.

Another key observation derived from the SCENIC analysis of these data is that cells in the MITF^high^ state also have high activity of STAT and IRF downstream targets. This is difficult to detect in bulk samples because of the complex mixture of malignant cells with tumor infiltrating lymphocytes (TIL) where STAT and IRF also play an important role ^79^. Here, we show that the MITF^high^ cells themselves have higher STAT activity than the MITF^low^ cells (we excluded all benign cells from the analysis, including immune cells). This has important consequences for the interpretation and prediction of resistance to immune therapy, because these cancer cells with high STAT and IRF activity are likely most sensitive to immunotherapy. Indeed, a recent study identified the JAK-STAT-IRF axis as driver for the expression of two major targets in immune therapy: PD-L1 and PD-L2; which results in an inhibition of the anti-tumor immune response on the one hand, but an increased response to anti-PD(L)1 immune therapies on the other ^80^.

In conclusion, in this work we provide a generally applicable method for the analysis of single-cell RNA-seq data, and for the first time exploit transcription factors and *cis*-regulatory sequences to guide the discovery of cellular states.

## Materials and methods

### SCENIC workflow

SCENIC is a workflow based on three new R/bioconductor packages: (1) GENIE3, to identify potential TF targets based on co-expression, (2) RcisTarget, to perform the TF-motif enrichment analysis and identify the direct targets (regulons), and (3) AUCell, to score the activity of regulons (or other gene sets) on single cells. We also provide GRNboost, implemented on Spark, as scalable alternative to build the co-expression network on bigger datasets (step 1).

The three R/bioconductor packages, and GRNboost, include detailed tutorials to facilitate their use within an automated SCENIC pipeline, as well as independent tools. Links to the tools, SCENIC code and tutorials are available at http://scenic.aertslab.org.

### GENIE3

GENIE3 is a method for inferring gene regulatory networks from gene expression data. In brief, it trains random forest models predicting the expression of each gene in the dataset, using as input the expression of the transcription factors. The different models are then used to derive weights for the transcription factors, measuring their respective relevance for the prediction of the expression of each target gene. The highest weights can be translated into TF-target regulatory links ^32^. GENIE3 is available in Python, Matlab and R. To allow for inclusion in SCENIC workflow, we optimized the previous R implementation of GENIE3. The core of this new implementation is now written in C –which makes it orders of magnitude faster–, it requires lower memory, and supports execution in parallel. GENIE3 was the top-performing method for network inference in the DREAM4 and DREAM5 challenges ^33^. The new package provides similar results in the DREAM challenge to previously existing implementations, but with improved speed. The comparison is available at the developer’s website: http://www.montefiore.ulg.ac.be/~huynh-thu/GENIE3.html.

The input to GENIE3 is an expression matrix. The preferred expression values are gene-summarized counts (which might or might not use unique molecular identifiers, UMI ^81^). Other measurements, such as counts or transcripts per million (TPM) and FPKM/RPKM are also accepted as input. However, note that the first network-inference step is based on co-expression, and some authors recommend avoiding within sample normalizations (i.e. TPM) for this task because they may induce artificial co-variation ^82^. To evaluate to what extent the normalization of the input matrix affects the output of SCENIC, we also ran SCENIC on Zeisel’s dataset after library-size normalization (using the standard pipeline from scran ^83^, which performs within-cluster size-factor normalization). The results are highly comparable, both in regards to resulting clusters/cell types (ARI between the cell types obtained from raw UMI counts or normalized counts: 0.90, ARI from normalized counts compared to the author’s cell types: 0.87), and the TFs identifying the groups (26 out of the 30 regulons highlighted in Figure 2b). Furthermore, during the course of this project we have applied GENIE3 to multiple datasets using UMI counts (e.g. mouse brain and oligodendrocytes), and TPM (e.g. human brain and melanoma) and both units provided reliable results.

The output of GENIE3 is a table with the genes, the potential regulators, and their “importance measure” (IM), which represents the weight that the transcription factor (input gene) has in the prediction of the target. We explored several ways to determine the threshold (e.g. looking at the rankings, distributions and outputs after pruning with RcisTarget), and finally opted for building multiple gene-sets of potential targets for each transcription factor: (a) setting several IM thresholds (IM > 0.001 and IM>0.005), (b) taking the 50 targets with highest IM for each TF, and (c) keeping only the top 5, 10 and 50 TFs for each target gene (then, split by TF). In all these cases, only the links with IM>0.001 were taken into account. Furthermore, each gene-set was then split into positive- and negative-correlated targets (i.e. Spearman correlation between the TF and the potential target) to separate likely activated and repressed targets. Finally, only the gene-sets (TF co-expression modules) with at least 20 genes were kept for the following step.

### GRNboost

GRNboost is based on the same concept as GENIE3: inferring regulators for each target gene purely from the gene expression matrix. However, it does so using the gradient boosting machines (GBM) ^72^ implementation from the XGBoost library ^84^. A GBM is an ensemble learning algorithm that uses boosting ^85^ as a strategy to combine weak learners, like shallow trees, into a strong one. This contrasts with random forest, the method used by GENIE3, which uses bagging (bootstrap aggregation) for model averaging to improve regression accuracy. GRNBoost uses gradient boosted stumps (regression trees of depth 1) ^86^ as the base learner. GRNBoost’s main contribution is casting this multiple regression approach into a Map/Reduce ^87^ framework based on Apache Spark ^52^. In GRNBoost, the core data entry is a tuple of a gene and a vector of TF expression values. Using a Spark RDD, GRNBoost first partitions the gene expression vectors over the nodes available in the compute cluster. Subsequently, it constructs a predictor matrix that contains the expression values for all candidate regulator genes. Using a Spark broadcast variable, the predictor matrix is broadcasted to the different compute partitions. In the map phase of the framework, GRNBoost iterates over the gene tuples (expression vector) and uses the predictor matrix to train the XGBoost regression models with the expression vectors as respective training labels. From the trained models, the strengths of the regulator-target relationships are extracted and emitted as a set of network edges. In the reduce phase, all sets of edges are combined into the final regulatory network.

The performance of GRNBoost and GENIE3 was compared on a workstation with 2 Intel Xeon E2696 V4 CPUs with in total 44 physical cores or 88 threads and 128 GB of 2133Ghz ECC memory. Large datasets and hence large predictor matrices cause the network inference to become memory-bound rather than CPU-bound. In order to comfortably fit the amount of memory required into the available 128 GB of memory, we decreased the number of partitions to 11, therefore having only a maximum of 11 predictor matrices in flight simultaneously. However, we increased the number of threads available to each individual XGBoost regression to 8, effectively using all available (88) threads in the workstation. GRNBoost is written in the Scala programming language and can be used as a software library or be submitted as a Spark job from the command line.

### RcisTarget

RcisTarget is a new R/Bioconductor implementation of the motif enrichment framework of i-cisTarget and iRegulon. RcisTarget identifies enriched transcription factor binding motifs and candidate transcription factors for a gene list. In brief, RcisTarget is based on two steps. First, it selects DNA motifs that are significantly over-represented in the surroundings of the transcription start site (TSS) of the genes in the gene-set. This is achieved by applying a recovery-based method on a database that contains genome-wide cross-species rankings for each motif. The motifs that are annotated to the corresponding TF and obtain a Normalized Enrichment Score (NES) > 3.0 are retained. Next, for each motif and gene-set, RcisTarget predicts candidate target genes (i.e. genes in the gene-set that are ranked above the leading edge). This method is based on the approach described by Aerts et al. ^88^ which is also implemented in <u>i-cisTarget</u> (web interface) ^89^ and <u>iRegulon</u> (Cytoscape plug-in) ^40^. Therefore, when using the same parameters and databases, RcisTarget provides the same results as i-cisTarget or iRegulon, benchmarked against other TFBS-enrichment tools in Janky et al. ^40^. More details about the method and its implementation in R are given in the package documentation.

To build the final regulons, we merge the predicted target genes of each TF-module that show enrichment of any motif of the given TF. To detect repression, it is theoretically possible to follow the same approach with the negative-correlated TF modules. However, in the datasets we analyzed, these modules were less numerous and showed very low motif enrichment, suggesting that these are lower quality modules. For this reason, we finally decided to exclude the detection of inhibition from the workflow, and continue only with the positive-correlated targets. The databases used for the analyses presented in this paper are the “18k motif collection” from iRegulon (gene-based motif rankings) for human and mouse. For each species, we used two gene-motif rankings (10kb around the TSS or 500bp upstream the TSS), which determine the search space around the transcription start site.

### AUCell

AUCell is a new method that allows identifying cells with active gene regulatory networks in singlecell RNA-seq data. The input to AUCell is a gene set or regulon, and the output the regulon “activity” (AUC) in each cell. In brief, AUCell’s scoring method is based on a recovery analysis, where the x-axis (Figure 1c) is the ranking of all genes based on expression level (genes with the same expression value, e.g. ‘0’, are randomly sorted); and the y-axis is the number of genes recovered from the input set. AUCell then uses the “Area Under the Curve” (AUC) to calculate whether a critical subset of the input gene set is enriched at the top of the ranking for each cell. In this way, the AUC represents the proportion of expressed genes in the signature and their relative expression value compared to the other genes within the cell. The output of this step is a matrix with the AUC score for each gene-set in each cell. We use either the AUC scores (across regulons) directly as continuous values to cluster single-cells, or we generate a binary matrix using a cutoff of the AUC score for each regulon. These cutoffs are determined either automatically, or manually adjusted by inspecting the distribution of the AUC scores. Some examples of AUC distributions are provided in Figure S1. The tutorial included in the package, also includes practical explanations and implications of each of the steps of the method.

### Cell clustering based on GRNs

The cell-regulon activity is summarized in a matrix in which the columns represent the cells and the rows the regulons. In the *binary* regulon activity matrix, the coordinates of the matrix that correspond to active regulons in a given cell will contain a “1”, and “0” otherwise. The equivalent matrix, containing the continuous AUC values for each cell-regulon, is normally referred to as the AUC activity matrix. Clustering of either of the regulon activity matrices reveals groups of regulons (jointly a network) that are recurrently active across a subset of cells. The binary activity matrix tends to highlight higher-order similarities across cells (and therefore, highly reduces batch effects and technical biases), on the other hand, the AUC matrix allows to observe more subtle changes. For visualization, we have mostly used t-SNEs (Rtsne package^90^, always tested consistency across several perplexity values and distance metrics/number of PCs), and heatmaps with hierarchical clustering (although the heatmap figures feature selected regulons, the t-SNEs are always run on the whole matrices). In the tutorials, we have also included several options to explore the results. For example, to detect most likely stable states (higher-density areas in the t-SNE), and to help identify key regulators, known cell properties (based on the dataset annotation) and GO terms (GO enrichment analysis of the genes in the cluster of regulons) that might be associated to the detected states.

### Data sources

#### Mouse cortex and hippocampus (Zeisel et al.)

The mouse brain dataset, published by Zeisel et al. ^16^, includes single-cell RNA-seq of 3005 cells from somatosensory cortex and hippocampus (CA1 region) of juvenile mice (21-31 days old). Most of the cells were sequenced after dissociation with no specific selection by markers or cell type (wild type CD-1 mice). In addition, the dataset also includes 116 cells selected by FACS from 5HT3a-BACEGFP transgenic mice (likely Htr3a interneurons). The expression matrix was downloaded from GEO (GSE60361). This matrix contains the UMI counts for 19972 genes across the 3005 cells that passed their quality controls (e.g. low quality cells and potential doublets). To run GENIE3, this matrix was filtered to keep the 13063 genes with more than 90 counts (which corresponds to 3 counts in 1% of cells) and detected in more than 30 cells (1% of cells). The rest of the SCENIC workflow was run as described in the previous section, leading to an activity matrix including 151 regulons. For the purpose of visualization, very sparse regulons can be filtered-out. For example, in Figure 2, we have plotted only regulons active in at least 1% of the cells and correlated with other regulons in the matrix (absolute correlation > 0.30). However, the downstream analyses include all the regulons.

#### Human neurons (Lake et al.)

The human neurons dataset, published by Lake et al. ^43^, includes single-cell RNA-seq data of 3083 neuronal cells from a normal brain (retrieved postmortem from a 51-year old female, from six different Brodmann areas: BA8, BA10, BA17, BA21, BA22, BA41/42). The expression matrix (available at the host laboratory webpage: http://genome-tech.ucsd.edu/ZhangLab/index.php/data/epigenomics-and-transcriptomics/sns/) contains expression values (in TPM) for 25122 genes in 4039 cells. Of these, only 3083 cells are retained after filtering out low mapping outliers and potential doublets. Repeated genes, mitochondrial genes and non protein coding genes were removed and the matrix was renormalized as log2(TPM+1). To run GENIE3, this matrix was filtered to keep the 14941 genes with more than 154 normalized counts (which corresponds to 5 normalized counts in 1% of cells) and detected in more than 31 cells (1% of cells). The rest of the SCENIC workflow was run as described in the previous section, except that the selected AUCell threshold was 0.20 instead of 0.03, leading to an activity matrix including 130 regulons.

#### Human brain (Darmanis et al.)

The human brain data set from Darmanis et al.^41^ provides scRNA-seq data from 466 cells from adult and fetal human brains. The fetal samples were taken from four different individuals at 16 to 18 weeks post-gestation. The adult brain samples were taken from healthy temporal lobe tissue (according to the test) from 8 different patients (21, 22, 37, 47, 50, 63 years old) during temporal lobectomy surgery for refractory epilepsy and hippocampal sclerosis. The expression profiles for 22085 genes in each cell (expressed as raw reads) were downloaded from GEO (GSE67835), merged into an expression matrix, and converted to logged CPM [log_10_(Reads per gene in a cell/Total reads in a cell)*1000000+1)]. Genes expressed in overall with less than 9.32 logged CPM counts (corresponding to at least 2 counts in 1% of the population) and expressed in less than 5 cells (1% of the population) were removed, resulting in an expression matrix with 14703 genes. The rest of the SCENIC workflow was run as described previously, resulting in a Regulon Activity Matrix with 259 regulons.

#### Mouse retina (Macosko et al.)

The dataset from Macosko et al. ^45^ contains scRNA-seq data of 44808 cells (after pruning singletons) obtained through Drop-seq from mouse retina (14 days post-natal). The expression matrix was obtained from GEO (GSE63472), while the cluster information was obtained from the host laboratory webpage (http://mccarrolllab.com/dropseq/). We used the normalized expression matrix [given as *log((UMI counts per gene in a cell/Total UMI counts in cell)*10000)+1)*].

In order to reduce the computational cost of the analysis, the dataset was down-sampled into a smaller set in which all the given cell types are represented. In mouse retina, the majority of the cells are rods ^91^, which according to the authors, in this data set correspond to more than 29000 cells. Since rods are the smallest cell type in mouse retina ^92^ and express fewer genes, they also contain higher levels of noise. In order to take a representative sample not overtaken by the rods content, Macosko et al. selected cells which express more than 900 genes. We used this same down-sampling approach, which resulted in a selection of 11020 cells, to build the gene regulatory network. Running GENIE3 (and RcisTarget) on the 12953 genes with more than 55.1 normalized counts (0.5 in 1% of the population) and detected in more than ~55 cells (0.5% of the population). This network was then evaluated on all the cells in the dataset, which led to an activity matrix including 123 regulons.

#### Embryonic mouse brain (10X Genomics)

The Chronium Megacell demonstration dataset contains 1,306,127 cells from cortex, hippocampus and subventricular zone of two E18 mice (strain: C57BL/6). We downloaded the expression matrix from the authors website (https://support.10xgenomics.com/single-cell-gene-expression/datasets/1Mneurons), which contains the expression data as 3 arrays in CSC (compressed sparse columnar) format compressed into a HDF5 file. Several subsets of this matrix were used to benchmark GRNboost (See GRNboost section).

#### Mouse oligodendrocytes (Marques et al.)

The oligodendrocytes data set from Marques et al. ^58^ contains scRNA-seq data of 5069 cells from the oligodendrocyte lineage. Cells were obtained from several different mouse strains and isolated from ten different regions of the anterior-posterior and dorsal-ventral axis of the mouse juvenile and adult CNS; including white and grey matter. The expression matrix, downloaded from GEO (GSE75330), provides the expression values in UMI counts for 23556 genes in those 5069 cells. Genes expressed in overall with less than 100 counts (corresponding to 2 counts in 1% of the population) or expressed in less than ~51 cells were filtered out for GENIE3 analysis, resulting in 11985 genes. The rest of the SCENIC workflow was run as described in the previous section, leading to an activity matrix including 128 regulons.

#### Oligodendroglioma (Tirosh et al.)

The oligodendroglioma data set from Tirosh et al.^76,75^ includes scRNA-seq expression profiles for 4347 cells from 6 untreated grade II oligodendroglioma tumors with either IDH1 or IDH2 mutation, and 1p/19q co-deletion. The expression data, given as log_2_(TPM+1), was downloaded from GEO (GSE70630). We only used the tumoral cells for the analysis. Most of the non-tumoral cells were removed from the data set by the authors based on CNV profile analysis. However, a total of 303 non-tumoral cells that lacked detectable CNVs were still included in the data set. We removed these non-tumoral cells from the expression matrix using hierarchical clustering based on the markers cited in the article (mature oligodendrocytes and microglia, respectively). Out of the 23686 genes in the expression matrix, we run GENIE3 on the genes expressed with more than 202 logged TPM counts (at least 5 logged TPM counts in 1% of the population) and detected in more than 40 cells (1% of the total data set), resulting in an expression matrix with 14728 genes and 4043 cells. The SCENIC pipeline was executed as previously described, resulting in a Regulon Activity Matrix with 159 regulons.

#### Melanoma (Tirosh et al.)

The melanoma dataset from Tirosh et al. ^55^ provides expression profiles of 23689 genes in 4645 cells from 19 melanoma tumors. These cells include both, malignant (melanoma cells), and non-malignant cells (e.g. immune cells). Here we analyze the 1252 melanoma cells (from 14 different tumors) that are labeled as malignant by the authors based on their CNV profiles. The expression matrix, as downloaded from GEO (GSE72056, on Apr 2016), is provided as logged TPM [log2(TPM/10+1)]. Therefore, for running GENIE3 we included the 14566 genes with more than 62.6 normalized counts per row (5 x 12.52 cells), that were detected (expression>0) in more than 12 cells (1%). In this way, the application of SCENIC on this dataset, lead to an activity matrix including 185 regulons.

### Gene and cell filtering

For gene filtering to run GENIE3, we applied a soft filter based on the total number of counts of the gene, and the number of cells in which it is detected. The first filter, the total number of reads per gene, is meant to remove genes that are most likely unreliable and provide only noise. The specific value depends on the dataset, for the ones used in this paper we set the thresholds at, for example, 3 UMI counts (slightly over the median of the non-zero values) multiplied by 1% of the number of cells in the dataset (e.g. in mouse brain: 3 UMI counts x 30 (1% of cells) = minimum 90 counts per gene). The second filter, the number of cells in which the gene is detected (e.g. >0 UMI, or >1 log2(TPM)), is to avoid that genes that are only expressed in one, or very few cells, gain a lot of weight if they happen to coincide in a given cell. In the workflow, we recommend to set a percentage lower than the smallest population of cells to be detected. For example, we initially considered setting this threshold at 5% (which might be ok for many datasets), but using this threshold with the mouse brain dataset, would have potentially left out many co-expression modules associated to the microglia cells, which, according to the authors, are approximately 3% of the total cells in the dataset. In this way, we finally required the genes to be detected in at least 1% of the cells.

### Method comparison for cell clustering

To determine whether the clustering based on gene regulatory network activity matches real cell types, we calculated sensitivity and specificity for the GRN assigned to the cells of the mouse brain (Figure 2c) according to the cell types assigned by Zeisel et al. For the other datasets, we compared the clustering (mainly t-SNE) based on the regulon activity matrices to the cell labels provided in the corresponding publications. For the comparison with clustering methods, we build on the benchmark presented in the SC3 publication ^27^, which also uses Zeisel et al., and provides the adjusted Rand index (ARI) on this dataset of 6 clustering methods commonly used for singlecell RNA-seq data. We extended the comparison with SEURAT by adding the results obtained with different resolution values.

### Method comparison for batch effects

The comparison of batch effect removal methods was performed using the Oligodendroglioma dataset by Tirosh et al. ^75^. SCENIC was run in the standard way (see Methods: Oligodendroglioma dataset), the t-SNE and diffusion plots were run on the full binary regulon activity matrix (the heatmap in Figure 4 illustrates selected stage-specific regulons). Combat ^59,60^ and Limma ^61,62^ were run to correct for “patient of origin” as source of batch effect (input matrix: 14728 genes and 4043 cells, same as GENIE3/SCENIC). Diffusion plots were done using the R/Bioconductor package destiny ^93,94^.

### Method comparison for cycling cells

Gene sets used to identify cycling cells are 46 sets related to the mitotic cell cycle, with at least 10 genes, retrieved from amiGO and cycleBase 1.0 and 2.0; Cycling cells are those that show consistent up-regulation of these genes, and are selected by hierarchical clustering on the matrix containing the z-scores for each gene-set. To compare methods, since most of the methods provide multiple clusters as output, for each method we selected the cluster with the biggest amount of CC cells. This cluster was then used to calculate the precision and recall.

**Combat** ^59,60^ and **Limma** ^61,62^ were run using the same logged and filtered matrix as in SCENIC. The tumor of origin was specified as the source of batch effect and hierarchical clustering (Ward’s method) was performed in the batch corrected matrix. The number of clusters was determined by the cutreeDynamic function of the dynamicTreeCut package ^87^. **GiniClust** ^20^ was run on the unlogged TPM matrix with the default parameters, which resulted in a matrix with 17843 genes and one single cluster. **PAGODA** ^13^ was run using the unlogged matrix (filter: 13976 genes with more than 100 TPM counts per row –at least 3 TPM counts in 1% of the population– and detected in more than 12 cells). The error models were generated using a k-nearest neighbor model fitting (minimum of 2 reads for the gene to be initially classified as a non-failed measurement, and at least 5 non-failed measurements per gene). The models were fitted based on 1/2 of the most similar cells (assuming that there will be two main subpopulations). These thresholds were established to avoid the impact of the TPM normalization of the counts. The variance normalization –which aims to normalize out technical bias and biological noise– was performed by trimming the 3 most extreme cells and limiting the maximum adjusted variance to 5. The rest of the PAGODA steps were run using default parameters, except: The evaluation of overdispersion of ‘de novo’ gene sets was performed with a trimming value of 7.1 extreme cells, and 50 as the number of clusters to be determined; threshold in the determination of top aspects a p-value: 0.01; distance threshold for the reduction of redundancy: 0.9. The optimal number of clusters in the data set was set to 10. **pcaReduce** ^96^ was run using the unlogged TPM matrix, with a similar filter as in SC3. The number of times to repeat the pcaReduce framework was set to 100 and the number of dimensions to start with was set to 30. Both merging methods (sampling based, S, and probability based, M) were tested. The number of clusters in the data set selected was 17 in both cases. **RaceID** ^17^ was run using an unlogged expression matrix filtered with the package’s function filterdata. The genes expressed with less than 8 TPM counts in at least 13 cells were discarded, resulting in 10371 genes. This matrix was normalized according to the RaceID procedure. The rest of the RaceID pipeline was run using default parameters. **SC3** ^14^ was run using the unlogged expression matrix. Default filters were applied, resulting in a matrix with 11645 genes. The number of estimated clusters was set to 21 (based on the output of the *sc3_estimate_k* function). **SEURAT** ^45^ was run using the unlogged TPM matrix and default parameters, correcting for tumor of origin as batch effect. The FindClusters function provided 8 clusters. **SIMLR** ^31^ was run with default paramaters using the unlogged expression matrix, with a similar filter to SC3. The number of clusters to be specified was set to 15. SINCERA ^26^ was run using the unlogged TPM matrix. A gene-by-gene per-cell z-score transformation was performed during the analysis, and hierarchical clustering was applied using correlation distances and average clustering. The number of clusters identified in the data set defined by the tool was 12. This threshold is set as the highest number of possible clusters that generates less than 1 singleton cluster. **SNNCliq** ^15^ was run using the scripts provided at http://hemberg-lab.github.io. The number of clusters found in the data set was 15. All these methods were run in R, except for SNNCliq, which is implemented in Python. Note that there are two tools related to cell cycle and single-cell RNA-seq datasets which have not been included in this analysis: (1) scLVM allows to correct for cofounding factors. These can be given in the form of a gene-set, for example GO gene-sets related to cell cycle, to correct for the cell cycle effect. However, to our knowledge, it does not provide an explicit score of the gene set on the cells. (2) Cyclone is a method to split cells according to their cell cycle stage. However, it assigns a cell cycle state to all the cells in the dataset, and thus, it is not useful for our purpose of identifying the cycling cells.

### Method comparison for TF-motif discovery

We compared SCENIC to an alternative approach to identify TFs potentially regulating cell states: Applying transcription factor motif enrichment analysis on genes differentially expressed between clusters (i.e. gene signature, or markers for a cell type). To do so, we started from the ‘gold standard’ for the mouse brain dataset: the cell type labels assigned by the authors, which are based on their own biclustering algorithm (Backspin) plus annotation based on markers to find the correspondence of each cluster to a given cell type. The ‘signatures’ for each cell type or cluster are defined based on four alternative approaches: (1) The genes assigned by Backspin to each cluster, (2) differentially expressed genes of each cluster versus all other cluster (One versus rest, OvR), (3) differentially expressed genes in each cluster versus any of the other clusters (One versus any other, OvAny), (4) highly variable genes across clusters (HVG). All the differential expression analyses, and the identification of HVG were run using EdgeR. We also run the best performing method, OvR, with an alternative differential expression tool, MAST, to confirm that the differential expression tool doesn’t have a major impact on the results. Homer and RcisTarget were then run on each of the gene-sets resulting from these contrasts: Backspin (pyramidal: 1960g, interneurons: 1126g, oligodendrocyte: 579g, microglia: 392g, astrocytes (only): 206g), OvR edger (interneurons: 574g, pyramidal: 1031g, oligodendrocytes: 859g, microglia: 1421g, endothelial_mural: 1541g, astrocytes_ependymal: 1121g), OvR MAST (pyramidal: 818g, interneurons: 688g, oligodendrocytes: 1057g, astrocytes_ependymal: 487g, microglia: 607g, endothelial_mural: 549g), OvAny (astrocytes_ependymal: 871g, endothelial_mural: 1019g, interneurons: 916g, microglia: 1060g, oligodendrocytes: 1013g, pyramidal: 816g), and 2013 highly variable genes (FDR<0.01, logFC>1).

Homer was run using the default parameters (promoter region: -300 to +50bp around TSS, motifs of length 8,10,12 and masking repeats). For the rest of the analysis we only took into account known motifs (ignored de-novo motifs), using the TF on the motif name as annotation to transcription factors. The equivalent analysis was also run with RcisTarget (the tool used by SCENIC), also using the default parameters with the two available databases (0 to - 500b, and - 10kbp to +10kbp around TSS) and only ‘direct annotation’ (TFs annotated to the motif in the original motif source). With both tools, we only took into account those TFs that are differentially expressed themselves, as this is also a standard approach to prioritize TFs and reduce false positives. For comparison with SCENIC, we used the results on the mouse brain presented in this paper (“diamond” shape in Figure 2f). Two versions of statistics are presented: One including all the TFs returned by SCENIC (including the sparse regulons), and one including only the “cell type” regulons, the regulons that are mostly specific to one of the final clusters. During the progress of this project, we updated the database of RcisTarget to contain about 2k more motifs, and developed GRNboost. Therefore, the figure includes the results for SCENIC using these new features (labelled as “DBv2” and “GRNboost”). As “true” TFs for the validation (Table S2), we took gene sets from mouse genome informatics (MGI) mammalial phenotype database (sample terms: abnormal [cell type] morphology/physiology, increased/decreased [cell type] number, abnormal brain morphology, abnormal blood-brain barrier function) and using the cell type as keyword (oligodendrocyte, astrocyte, interneuron, pyramidal, neuron, microglia, brain endothelial), and from the gene ontology (e.g. GO:0014013 Regulation of gliogenesis, GO:0022008 Neurogenesis,…). Finally, the precision and recall for each method were calculated according to the TFs identified across all the cell types (e.g. joining all cell types, not cell-type specific), since multiple cell-type specific TFs known from literature were only available in generic terms (e.g. “brain”, “gliogenesis”), not in the cell specific annotations.

### Cross-species network comparisons

SCENIC was run independently for each of the three datasets used for the GRN comparison: Zeisel et al. (mouse brain cells), and Lake et al. (human neurons nuclei) and Darmanis et al. (human brain cells). To compare the networks across species, the genes in the human regulons were converted into the homologous mouse genes using Biomart (through biomaRt R package ^97^), and vice versa (the mouse regulons into human genes). For the cross-species cell clustering (Figure 3c), the genes in the mouse expression matrix were converted into the homologous human genes, and merged with Darmanis’ expression matrix by row (only genes available in both matrices are kept). The 259 human regulons from Darmanis’ dataset, and the human homologs of the mouse regulons were evaluated on this merged matrix to obtain the binary regulon activity containing 410 regulons. The cells were clustered based on the binary activity matrix using Ward’s hierarchical clustering with Spearman’s distance. Similar results were obtained for the reverse approach (converting the expression matrix into mouse genes, to evaluate the mouse regulons). In order to provide an alternative approach based only on expression (Figure S9), we also generated a merged expression matrix. Since the merged data sets use different measurement units (CPM in human and UMI in mouse), each matrix was Z-score normalized by gene before merging.

### Gene Ontology

To identify enriched GO terms or pathways associated to the states identified, we performed functional enrichment analysis on the union of regulons associated to each state (i.e. for the synaptic oligodendrocyte group, the union of targets of Hoxa7, Hoxa9, Hoxb7 and Hoxb9). The analyses were performed mainly through Mouse Mine ^98^ (although we also checked DAVID ^99,100^, Enrichr ^101,102^, which provided similar results).

### Differential expression analysis

Differential expression between clusters of single cells was performed using MAST ^8^. Genes expressed with less than 9 counts in the population were excluded. Differentially expressed genes were evaluated according to their log fold change and adjusted p-values.

### Immunohistochemistry of melanoma biopsies

Immunohistochemistry was performed on formalin-fixed, paraffin-embedded melanoma samples on the Leica BOND-MAX^™^ automatic immunostainer (Leica Microsystems). Antigen retrieval was performed onboard using a citrate-based (Bond Epitope Retrieval Solution 1, pH 6.0; Leica) or a EDTA-based buffer (Bond Epitope Retrieval Solution 2, pH 9.0; Leica) according to the manufacturer’s instructions. The antibodies were used for melanA (IR633 from DAKO, initially diluted at RTU, but further diluted 1:2 for better contrast; antigen retrieval: pH9.0), EPHA2 (#6997 from Cell Signaling Technology diluted at 1/50; pH9.0), ZEB1 (sc-25388 from Santa Cruz Biotechnology diluted at 1/200; pH9.0), NFATC2 (#5861 from Cell Signaling Technology diluted at 1/5000; pH6.0) and NFIB (HPA003956 from Sigma Aldrich diluted at 1/250; pH6.0). Alkaline phosphatase activity was detected with Bond Polymer Refine Red Detection (Leica) as substrate, resulting in a pink/red immunoreactivity. To help identification of melanoma cells in sentinel lymph nodes, double immunohistochemical staining with melanA were performed with sequential development of peroxidase and alkaline phosphatase with Bond Polymer Refine Detection and Bond Polymer Refine Red Detection (Leica Microsystems, Wetzlar, Germany), respectively, resulting in contrasting dark brown (marker) and pink/red immunoreactivities (melanA).

### Knock-down of NFATC2 in melanoma cell culture

A375 cell line was selected based on expression of NFATC2, NFIB (Figure 5l), and SOX10 across 59 melanoma cell lines from the COSMIC Cancer Cell lines Project^103^. A375 cells were obtained from the ATCC and were cultured in Dulbecco’s Modified Eagle’s Medium with high glucose and glutamax (ThermoFisher Scientific), supplemented with 10% fetal bovine serum (Lonza) and penicillin-streptomycin (ThermoFisher Scientific). Knockdown of NFATC2 was performed using the ON-TARGETplus NFATC2 siRNA SMARTpool (Dharmacon) at a final concentration of 40nM in opti-MEM medium (ThermoFisher Scientific). Total RNA was extracted 72 hours after knockdown, using the innuPREP RNA mini kit (Analytik Jena), according to the manufacturer’s instructions. Quality checks were performed using the Bioanalyzer 1,000 DNA chip (Agilent) after which libraries were constructed: After total RNA purification, mRNA was enriched using the Dynabeads mRNA purification kit (Invitrogen). To make cDNA, 1 μl of oligo(dT) primers (500ng/μl; Ambion) and 1 μl of 10 mM dNTP (Promega) was added to 10 μl of polyA-selected mRNA; incubated at 65°C for 5 min and placed on ice. First-strand cDNA synthesis was performed by adding 4 μl of first strand buffer (Invitrogen), 2 μl of 100 mM DTT (Invitrogen) and 1 μl of Superscript II (Invitrogen) and incubating the mix at 42°C for 50 min, then 70°C for 15 min. The second strand of cDNA was filled in by adding 35 μl of water, 15 μl of 5x second strand buffer (Invitrogen), 1.5 μl of 10 mM dNTP, 0.5 μl of 10 U/μl E Coli DNA ligase (Bioke), 2 μl of 10 U/μl E Coli DNA polymerase I (Bioke), 1 μl of 2 U/μl E Coli RNaseH and then incubating at 16°C for 2 hours. The cDNA was purified on a MinElute column (Qiagen) and eluted in 15 μl EB buffer. To incorporate sequencing adapters, we combined the purified cDNA with 4 μl of Nextera TD buffer (Illumina) and 1 μl of Nextera Tn5 enzyme (Illumina) on ice and incubated at 55°C for 5 min. The tagmented cDNA was purified again on a MinElute column and eluted in 20 μl EB buffer. To PCR amplify the fragments, we added 25 μl of NEBnext PCR master mix (Bioke), 5 μl of Nextera primer mix and incubated at 72°C for 5 min, then at 98°C for 30 sec, followed by 15 cycles of 98°C for 10 sec, 63°C for 30 sec and 72°C for 3 min. We purified the PCR amplicons with 55 μl AMPure beads (Analis).

Final libraries were pooled and sequenced on a NextSeq 500 and HiSeq 4000 (Illumina). Raw Fastq files of the same sample were merged and adapter sequences were removed using fastq-mcf. RNA-seq reads were mapped to the genome (hg19) using STAR (v2.5.1b) and reads with high mapping quality (Q4) were selected using SAMtools (v1.4). Read counts per gene were obtained from the aligned reads using htseq-count. The Bioconductor/R package DESeq2 was used for normalization and differential gene expression analysis. Log2FoldChange values were used for ranking the genes, and downstream GOrilla and GSEA analysis, as previously described.

### Software packages, code and data availability

SCENIC workflow and updated links to all packages and tutorials: http://scenic.aertslab.org. Four new software packages have been developed within the context of this project. Current versions of the packages are available at: GENIE3: https://github.com/aertslab/GENIE3; RcisTarget: https://github.com/aertslab/RcisTarget (The motif databases required to perform analyses are available as separate data packages); AUCell: https://github.com/aertslab/AUCell; GRNboost: https://github.com/aertslab/GRNBoost. The scripts used for all analyses (including the generation of the figures) are provided on the SCENIC website.

The NFATC2 knock-down RNA-seq data have been deposited in NCBI’s Gene Expression Omnibus ^104^ and are accessible through GEO Series accession number <u>GSE99466</u>.

## Acknowledgements

This work is funded by The Research Foundation - Flanders (FWO, www.fwo.be; grants G.0640.13 and G.0791.14 to SA), Special Research Fund (BOF) KU Leuven (http://www.kuleuven.be/research/funding/bof; grant PF/10/016 and OT/13/103 to SA), Foundation Against Cancer (http://www.cancer.be; 2012-F2 to SA) and ERC Consolidator Grant (724226_cis-CONTROL to SA). SAS is funded by a PDM Postdoctoral Fellowship from the KU Leuven. ZKA and JW are funded by postdoctoral research felloships from Kom op Tegen Kanker. HI has a PhD Fellowships from the agency for Innovation by Science and Technology (IWT, www.iwt.be). The funders had no role in study design, data collection and analysis, decision to publish, or preparation of the manuscript. TM would like to thank Jaak Simm for helpful comments and suggestions regarding gradient boosting.

## List of supplementary figures

Figure S1. AUCell score distributions for multiple gene-sets

Figure S2. AUCell scores of known cell signatures on the mouse brain dataset

Figure S3. Comparison of the regulon activity and the TF expression alone

Figure S4. Association of brain networks with Alzheimer’s disease neurodegeneration

Figure S5. SCENIC is robust to downsampling of cells and sparse expression matrices

Figure S6. Benchmarking SCENIC

Figure S7. Binary regulon activity for the human neurons nuclei

Figure S8. Interneuron subtypes on the human single-nuclei data set

Figure S9. Expression-based clustering of mouse and human brain cells

Figure S10. Analysis of >40000 single cells

Figure S11. GRNboost: Comparison of performance with GENIE3

Figure S12. GRNboost: Evaluation of stability across multiple runs

Figure S13. Validation of the cell cycle cells and comparison with other methods

Figure S14. Oligodendrocyte differentiation is driven by discrete changes in gene regulatory networks

Figure S15. Comparison of TF expression and regulon activity

Figure S16. Expression of known melanoma markers

Figure S17. Comparison of melanoma bulk signatures with single-cell states

Figure S18. Immunohistochemistry of NFATC2, NFIB and ZEB1 on human melanomas

## List of supplementary tables

Table S1. Excel file with RNA-seq results of NFATC2 knock-down versus control

Table S2. Transcription factors used for validation in the TF-motif discovery comparisons

